# Lipidomic profiling reveals concerted temporal patterns of functionally related lipids in *Aedes aegypti* females following blood feeding

**DOI:** 10.1101/2022.05.18.492402

**Authors:** Meng-Jia Lau, Shuai Nie, Qiong Yang, Lawrence G. Harshman, Cungui Mao, Nicholas A. Williamson, Ary A. Hoffmann

## Abstract

We conducted a lipidomic analysis of the whole body of female *Aedes aegypti* mosquitoes at different time points over the course of feeding and reproduction. There were temporal biphasic increases of more than 80% of lipids identified at the time of feeding and from 16 h to 30 h post blood meal. During these two increases, the abundance of many lipids dropped while body weight remained stable, probably reflecting blood lipid digestion and the synthesis of vitellogenin in this period. A concerted temporal pattern was particularly strong at the second peak for membrane and signalling lipids such as PE, PI, CL, HexCer and LPA. Lyso-glycerophospholipids showed three distinct change patterns which are functionally related: LPE and LPC, which are membrane lipids, showed little change; LPA, a signalling lipid, showed a dramatic increase from 16 to 30 h PBM; LPI, a bioactive lipid, and both LPG and LPS which are bacterial membrane lipids, showed one significant increase from the time of feeding to 16 hours post blood meal. The result of our study on the anautogenous insect *Ae. aegypti* point to specific lipids likely to be important in the reproductive process with a role in the formation and growth of ovarian follicles.

## Introduction

The yellow fever mosquito, *Aedes aegypti*, is one of the major vectors that transmits arbovirus diseases in tropical and subtropical areas, such as dengue and Zika [1–3]. *Ae. aegypti* females are anautogenous: they must ingest human/vertebrate blood to obtain unique nutrients to produce their eggs, and diseases are transmitted through the blood ingestion process [4,5].

Reproduction in mosquito females is generally regulated through the provision of nutrition and the modulation of hormones [6]. Essential nutrients such as amino acids and cholesterol mainly come from human blood [7,8], and supports gene products involved in yolk development [9]. The source and the quality of blood that females are fed can directly impact their fertility [10–13], though their nutritional components such as amino acids and cholesterol are considered similar [14,15]. Apart from these changes of amino acids, cholesterol and yolk protein, the vittellogenic cycle is also expected to involve lipid metabolism; lipids not only act as major energy resources in supporting egg development [16] and contribute 35% of ovarian dry weight [17,18], but also exhibit a variety of functions in biological system [19,20]. The development of ovaries can be separated into the pre-vitellogenic (before blood meal, BBM) and vitellogenic (post blood meal, PBM) stages depending on the timing of the blood meal. In the pre-vitellogenic stages, lipids are reserved in the fat body [16], which is essential to support *Ae. aegypti* reproduction [21], particularly when a sugar source is compromised. Starving female mosquitoes before or after their blood feeding will not reduce their fecundity but will significantly reduce the amount of lipids in mothers after laying eggs [22], suggesting a trade-off between reproduction and female survival.

During the vitellogenic stages, consumption of blood meal triggers insulin-like peptide 3 (ILP3) from the brain to stimulate oogenesis [23], which represents the insulin signalling pathway [24] The secretion of 20-hydroxyecdysone then increases the expression of vitellogenin genes in the fat body to support yolk protein accumulation and ovarian follicle development [25], which is stimulated by gonadotropin [26], and lipids are transferred from the fat body to follicles [16]. In the meantime, *Ae. aegypti* females also digest and absorb essential amino acids from human blood, where the concentration of amino acids increasing in haemolymph acts as a signal to activate egg development [27] through the TOR pathway [7]. In this study, we used liquid chromatography coupled with the mass spectrometry (LCMS)-based lipidomic method and a comprehensive lipid standard mixture covering near all major lipid groups to identify and analyse lipid components of whole female individuals before the blood meal and post blood meal. Our study aims to understand lipid metabolism in *Ae. aegypti* during the reproductive process, and how lipids with different functions act to support the development of eggs.

## Material and methods

### Sample preparation

*Aedes aegypti* mosquitoes in this experiment were collected from *Wolbachia*-uninfected populations from Cairns, QLD, Australia and maintained in laboratory conditions. Samples were prepared by hatching *Ae. aegypti* eggs stored in an incubator (PG50 Plant Growth Chambers, Labec Laboratory Equipment, Marrickville, NSW, Australia) at 22-30 ± 1 °C, with a 12 h : 12 h photoperiod. Larvae were reared in reverse osmosis (RO) water with yeast and fed with Hikari Sinking Wafers fish food tablets until adult. All female mosquitoes were blood fed by the same volunteer’s forearm. The first blood meal was provided to adult females at 4 ± 1 days post-emergence. Females were isolated individually and observed for seven days to discard those that laid small numbers of eggs (< 50). At the seventh day, a second blood meal was provided. Non-engorged individuals were removed from the experiment. Samples were then stored as different groups in their second vitellogenic cycle before blood meal (BBM) and at different time post blood meal (PBM): 0 h PBM (30 mins after blood meal), 4 h PBM, 16 h PBM, 30 h PBM, 72 h PBM, and 120 h PBM. Sugar and oviposition cups were provided during the entire period of the experiment.

### Female mosquito measurement

Apart from analysis of the lipidome, we measured the weight of different female individuals (four to seven) at each time point from the additional collected samples using a Sartorius Analytical balance BP 210 D (Sartorius, Gottigen, Germany, Readability: 0.01 mg). After weighing, eight mosquitoes were randomly selected from all time points and their wing lengths were measured to indicate female body size [28]. To further understand the development of female ovaries, we hatched batches of mosquito eggs and took photos of ovaries using an NIS Elements BR imaging microscope (Nikon Instruments, Tokyo, Japan) every day PBM.

### Lipid extraction

For each group, we extracted the whole body of six mosquito females, keeping individuals separate. We added 10 μL of Mouse SPLASH® LIPIDOMIX® Mass Spec Standard (cat. 330707, Avanti) after 1 in 4 dilution to each sample prior the extraction and then homogenised samples in 200 μL ice-cold 60% methanol (MeOH) containing 0.01% (w/v) butylated hydroxytoluene (BHT) by using TissueLyserII (Qiagen, Hilden, Germany) at 25 Hertz for one minute and incubating for 20 minutes in an ultrasonic bath (Soniclean Pty Ltd., Thebarton, Australia). Following the method previously reported by Lydic, *et al*. [29], 120 μL of MilliQ water, 420 μl of MeOH with 0.01% (w/v) BHT, and 260 μL of chloroform (CHCl_3_) were added to each tube and vortexed thoroughly before incubating in a Bioshake iQ (Quantifoil Instrument GmbH, Germany) setting at 1,400 rpm for 30 minutes and centrifuged at 14,000 rpm for 15 minutes. Supernatant containing lipids was transferred to a new 2 mL tube. The remaining sediments were re-extracted with 100 μL of MilliQ water and 400 μL of CHCl_3_ : MeOH (1:2, v:v) containing 0.01% (w/v) BHT as described above. Supernatants from the first and second extraction were pooled in the same 2 mL tube and then dried by evaporation under vacuum using a GeneVac miVac sample concentrator (SP Scientific, Warminster, PA, USA). Before HPLC, dried lipid extracts were resuspended in 50 μL of CHCl_3_ : MeOH (1:9, v:v, containing 0.01% BHT).

### Mass spectrometry analyses

Samples were analysed by ultrahigh performance liquid chromatography (UHPLC) coupled to tandem mass spectrometry (MS/MS) employing a Vanquish UHPLC linked to an Orbitrap Fusion Lumos mass spectrometer (Thermo Fisher Scientific), with separate runs in positive and negative ion polarity. Solvent A was 6/4 (v/v) acetonitrile/water with 10 mM ammonium acetate and 5 μM medronic acid and solvent B was 9/1 (v/v) isopropanol/acetonitrile with 10 mM ammonium acetate. Each sample was injected into an RRHD Eclipse Plus C18 column (2.1 × 100 mm, 1.8 μm, Agilent Technologies) at 50°C at a flow rate of 350 μl/min for 3 min using 3% solvent B. During separation, the percentage of solvent B was increased from 3% to 70% in 5 min, from 70% to 99% in 16 min, from 99% to 99% in 3 min, from 99% to 3% in 0.1 min and maintained at 3% for 3.9 min.

All experiments were performed using a Heated Electrospray Ionization source. The spray voltages were 3.5 kV in positive ionisation mode and 3.0 kV in negative ionisation mode. In both polarities, the flow rates of sheath, auxiliary and sweep gases were 20 and 6 and 1 ‘arbitrary’ unit(s), respectively. The ion transfer tube and vaporizer temperatures were maintained at 350 °C and 400 °C, respectively, and the ion funnel RF level was set at 50%. In both positive and negative ionisation modes from 3 to 24 min, top speed data-dependent scan with a cycle time of 1 s was used. Within each cycle, a full-scan MS spectrum was acquired firstly in the Orbitrap at a mass resolving power of 120,000 (at *m/z* 200) across a *m/z* range of 300–2000 using quadrupole isolation, an automatic gain control (AGC) target of 4e5 and a maximum injection time of 50 milliseconds, followed by higher-energy collisional dissociation (HCD)-MS/MS at a mass resolving power of 15,000 (at *m/z* 200), a normalised collision energy (NCE) of 27% at positive mode and 30% at negative mode, an *m/z* isolation window of 1, a maximum injection time of 22 milliseconds and an AGC target of 5e4. For the improved structural characterisation of glycerophosphocholine (PC) in positive mode, a data-dependent product ion (*m/z* 184.0733)-triggered collision-induced dissociation (CID)-MS/MS scan was performed in the cycle using a q-value of 0.25 and a NCE of 30%, with other settings being the same as that for HCD-MS/MS. For the improved structural characterisation of triacylglycerol (TG) lipid in positive mode, the fatty acid + NH_3_ neutral loss product ions observed by HCD-MS/MS were used to trigger the acquisition of the top-3 data-dependent CID-MS3 scans in the cycle using a q-value of 0.25 and a NCE of 30%, with other settings being the same as that for HCD-MS/MS.

### Lipid identification and quantification

LC-MS/MS data was searched through MS Dial 4.80 [30,31]. The mass accuracy settings are 0.005 Da and 0.025 Da for MS1 and MS2. The minimum peak height is 50000 and mass slice width is 0.05 Da. The identification score cut off is 80%. Post identification was done with a text file containing name and *m/z* of each standard in Mouse SPLASH® LIPIDOMIX® Mass Spec Standard. In positive mode, [M+H]^+^ and [M+NH_4_]^+^ were selected as ion forms. In negative mode, [M-H]^-^ were selected as ion forms. All lipid classes available were selected for the search. PC, LPC, DG, TG, CE, SM were identified and quantified at positive mode while PE, LPE, PS, LPS, PG, LPG, PI, LPI, PA, LPA, Cer, CL were identified and quantified at negative mode. The retention time tolerance for alignment is 0.1 min. Lipids with maximum intensity less than 5-fold of average intensity in blank were removed. All other settings were default. All lipid LC-MS features were manually inspected and re-integrated when needed. We also removed four other types of lipids: (1) lipids with only sum composition except PC and SM; (2) lipids identified due to peak tailing; (3) retention time outliers within each lipid class; and (4) LPA and PA artifacts generated by in-source fragmentation of LPS and PS. The shorthand notation used for lipid classification and structural representation follows the nomenclature proposed previously [32].

Quantification of lipid species was achieved by comparison of the LC peak areas of identified lipids against peak areas and quantities of the corresponding internal lipid standards in the same or similar lipid class (Table 1), and the concentrations of all lipids were reported at nmol/individual. Given that the commercially available stable isotope-labelled lipid standards are limited, some of the identified lipids were normalised against a standard from a different class or sub-class, and no attempts were made to quantitatively correct for different ESI responses of individual lipids due to concentration, acyl chain length, degree of unsaturation, or matrix effects caused by differences in chromatographic retention times compared with the relevant standards.

**Table 1.** List of lipid standards, adduct type and number of lipids that were identified in each group.

| Name (Abbreviation) | Standard | Adduct type | Number of lipids identified |
| --- | --- | --- | --- |
| Diacylglycerol (DG) | 15:0-18:1(d7) DAG | $[M+NH_4]^+$ | 14 |
| Triglycerides (TG) | 15:0-18:1(d7)-15:0 TAG | $[M+NH_4]^+$ | 41 |
| Phosphatidylethanolamines (PE) | 15:0-18:1(d7) PE | $[M-H]^-$ | 43 |
| Ether-linked PE (EtherPE) | 15:0-18:1(d7) PE | $[M-H]^-$ | 18 |
| Oxidized PE (OxPE) | 15:0-18:1(d7) PE | $[M-H]^-$ | 8 |
| Phosphatidylinositol (PI) | 15:0-18:1(d7) PI | $[M-H]^-$ | 33 |
| Ether-linked PI (EtherPI) | 15:0-18:1(d7) PI | $[M-H]^-$ | 3 |
| Phosphatidylserine (PS) | 15:0-18:1(d7) PS | $[M-H]^-$ | 10 |
| Phosphatidic acids (PA) | 15:0_18:1(d7) PA | $[M-H]^-$ | 5 |
| Phosphatidylglycerol (PG) | 15:0_18:1(d7) PG | $[M-H]^-$ | 24 |
| Phosphatidylcholines (PC) | 15:0_18:1(d7) PC | $[M+H]^+$ | 52 |
| Ether-linked PC (EtherPC) | 15:0_18:1(d7) PC | $[M+H]^+$ | 7 |
| Lyso-PE (LPE) | 18:1(d7) Lyso PE | $[M-H]^-$ | 13 |
| Ether-linked lyso-PE (EtherLPE) | 18:1(d7) Lyso PE | $[M-H]^-$ | 2 |
| Lyso-PI (LPI) | 15:0-18:1(d7) PI | $[M-H]^-$ | 7 |
| Lyso-PS (LPS) | 15:0-18:1(d7) PS | $[M-H]^-$ | 6 |
| Lyso-PA (LPA) | 15:0_18:1(d7) PA | $[M-H]^-$ | 4 |
| Lyso-PG (LPG) | 15:0_18:1(d7) PG | $[M-H]^-$ | 7 |
| Lyso-PC (LPC) | 18:1(d7) Lyso PC | $[M+H]^+$ | 9 |
| Cardiolipin (CL) | 15:0_18:1(d7) PG | $[M-H]^-$ | 66 |
| Carnitine (CAR) | 15:0_18:1(d7) PC | $[M+H]^+$ | 15 |
| Sphingomyelin (SM) | d18:1-18:1(d9) SM | $[M+H]^+$ | 26 |
| Ceramides (Cer) | d18:1-18:1(d9) SM | $[M-H]^-$ | 23 |
| Hexosylceramides (HexCer) | d18:1-18:1(d9) SM | $[M-H]^-$ | 4 |
| Sulfatides (SHexCer) | d18:1-18:1(d9) SM | $[M-H]^-$ | 16 |

### Data analysis

All data were analysed through R v. 3.6.0. Female weight was analysed by ANOVA across time followed by posthoc Tukey HSB tests. The concentrations of different lipid groups (nmol/individual) were used as variables for principal component analysis (PCA). For specific comparisons between treatments for particular lipids, concentration was natural log-transformed before undertaking paired t-tests, and p values were corrected for lipid number considered in a comparison by the Benjamin-Hochberg method.

## Results

### Female body weight and wing length measurement

*Aedes aegypti* female mosquitoes were stored for lipidomic study in their second vitellogenic cycle before their blood meal (BBM) and at different time post blood meal (PBM): (0 h (30 mins after blood meal), 4 h, 16 h, 30 h, 72 h, and 120 h). We first weighed female mosquitoes from samples collected at each time point (Figure 1), with the difference between BBM and 0 h PBM representing the amount of blood a female engorged. Sampling time had a significant effect on female weight (ANOVA, F_7,41_ = 13.06, P < 0.001). The weights of engorged female mosquitoes were approximately double their weights BBM. The females then started to digest blood, and their weight dropped significantly from 0 h PBM to 4 h PBM, thereafter stabilising until 30 h PBM, and then dropping again further to the BBM level at 72 h PBM; this change in weight reflects the process of blood digestion, which takes around 32 hours after females are engorged [33,34]. The appearance of female ovaries was captured every day under a microscope (Supplementary Material 1). We also calculated the coefficient of variation (CV, CV = (Standard Deviation/Mean) × 100) for female wing length for the eight samples we measured (mean ± se = 2.07 ± 0.06 mm, CV = 8.71) to indicate the spread of mosquito body sizes in this study.

**Figure 1.**
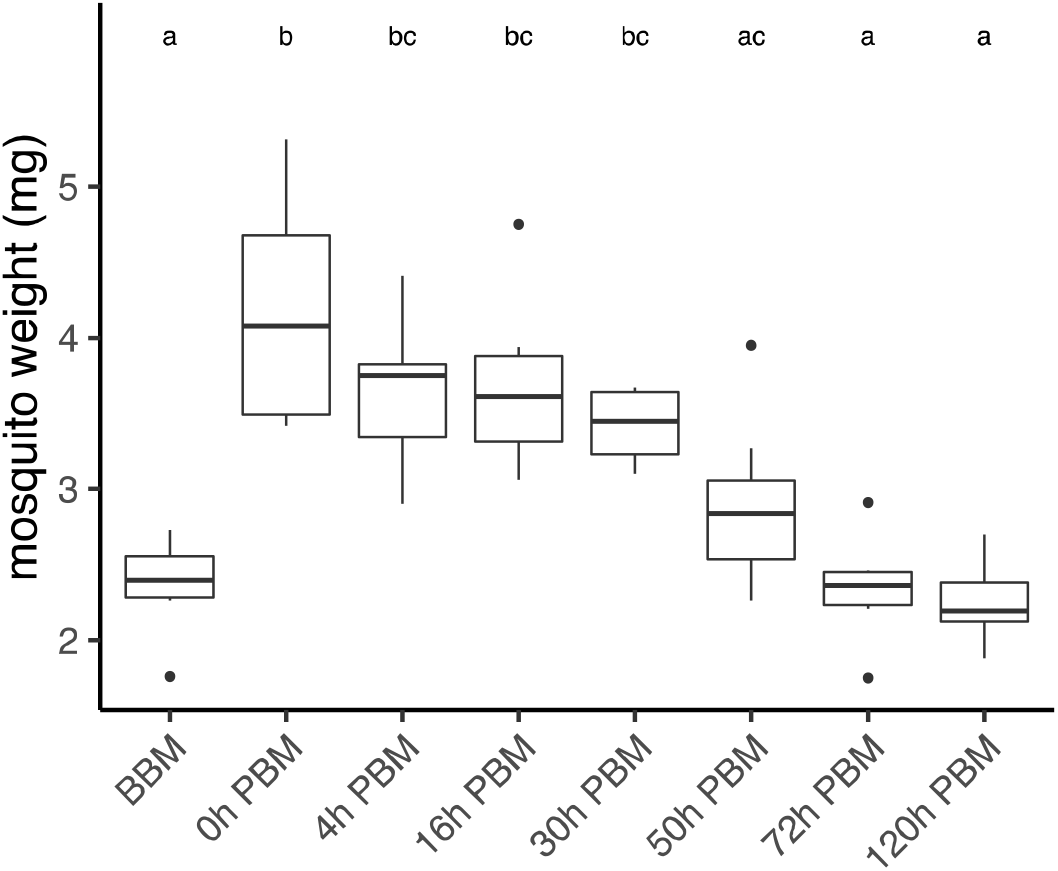
Weights of *Aedes aegypti* females across different time points over the course of feeding and reproduction. Weights were measured at eight time points before blood meal (BBM) and post blood meal (PBM). The same letter at the top of the boxplots represents groups that are not significantly different following Tukey HSD post-hoc tests.

### Lipids identification and their changes during female reproductive process

We measured lipidomic changes averaged to individual female BBM and at different time points PBM, covering the entire vitellogenic cycle. In total, we identified 456 lipids (Table 1) and divided them into six major groups: glycerolipids, diacyl-glycerophospholipids, lyso-glycerophospholipids, Cardiolipin (CL), Carnitine (CAR) and sphingolipids. The most abundant lipid group in female mosquitoes was the neutral glycerolipids, with 55 molecules making up approximately 80% of the total lipid amount. Glycerophospholipids accounted for 55% of the molecules identified in this class, with 193 diacyl-glycerophospholipids and 48 lyso-glycerophospholipids. We also identified 66 CL, 15 CAR and 69 sphingolipids.

In order to obtain an overview of lipidomic changes during the reproductive process, we first undertook a principal component analysis (PCA) to describe overall patterns (Figure 2). This indicated that lipid components of female mosquitoes went through two main changes: the first occurred immediately after a blood meal (0 h PBM), and the second change occurred between 16 and 30 h PBM. Further analysis showed that the amounts of most lipids increased at 0 h and 16 h PBM respectively (Figure 3), except for lyso-glycerophospholipids, in which case the total amount remained relatively stable during the reproduction process. These results are also reflected in PCA analyses on specific classes of lipids (Figure 4); for most classes the first PC explained more than half of the variation, with the exception of the lyso-glycerophospholipids where it only explained 31.8% of variation and there was no obvious change directly after blood feeding. However there was a shift at a later time point for this group, indicating that most changes in lyso-glycerophospholipid reflect mosquito metabolism rather than direct blood ingestion.

**Figure 2.**
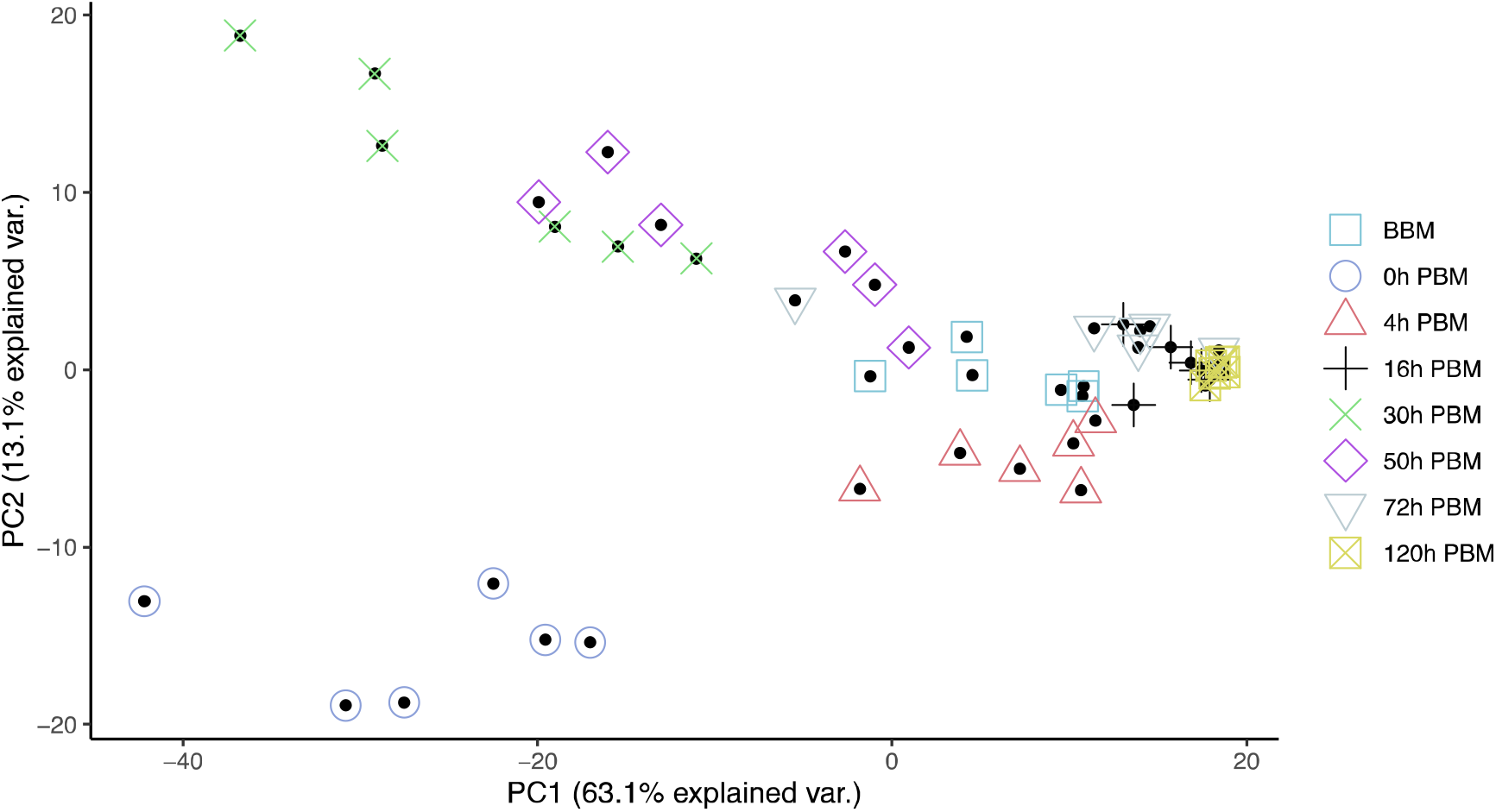
Principal Component Analysis suggested lipidome differences of *Aedes aegypti* females at different time points over the course of feeding and reproduction. Colour and shape symbols correspond to the eight time points before blood meal (BBM) and post blood meal (PBM). The first two principal axes explained 76 % of the variance.

**Figure 3.**
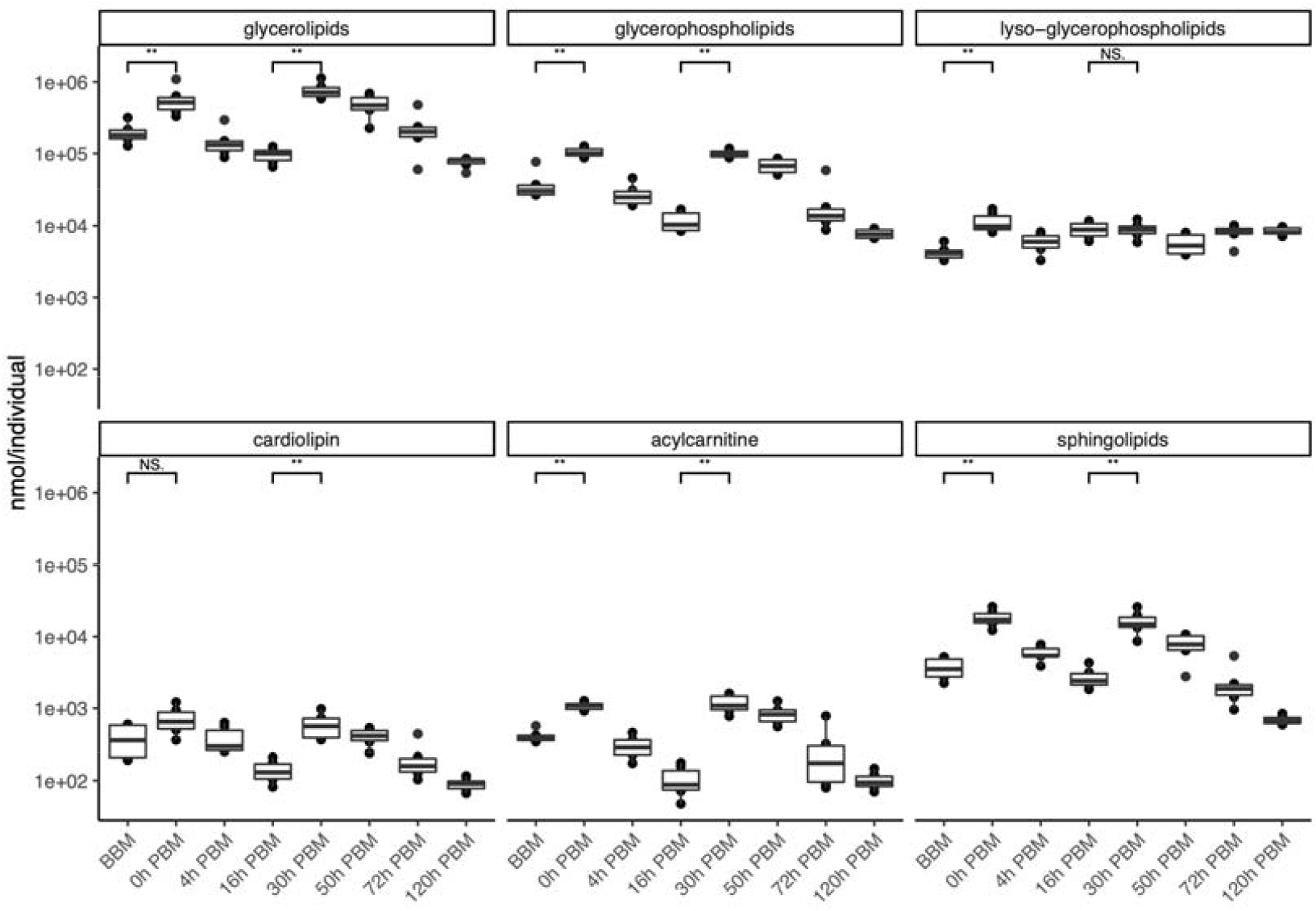
Changes of lipid amounts in *Aedes aegypti* females over the course of feeding and reproduction. The total amount (nmol/individual) of glycerolipids, diacyl-glycerophospholipids, lyso-glycerophospholipids, cardiolipins, carnitines and sphingolipids were measured at eight time points before blood meal (BBM) and post blood meal (PBM).

**Figure 4.**
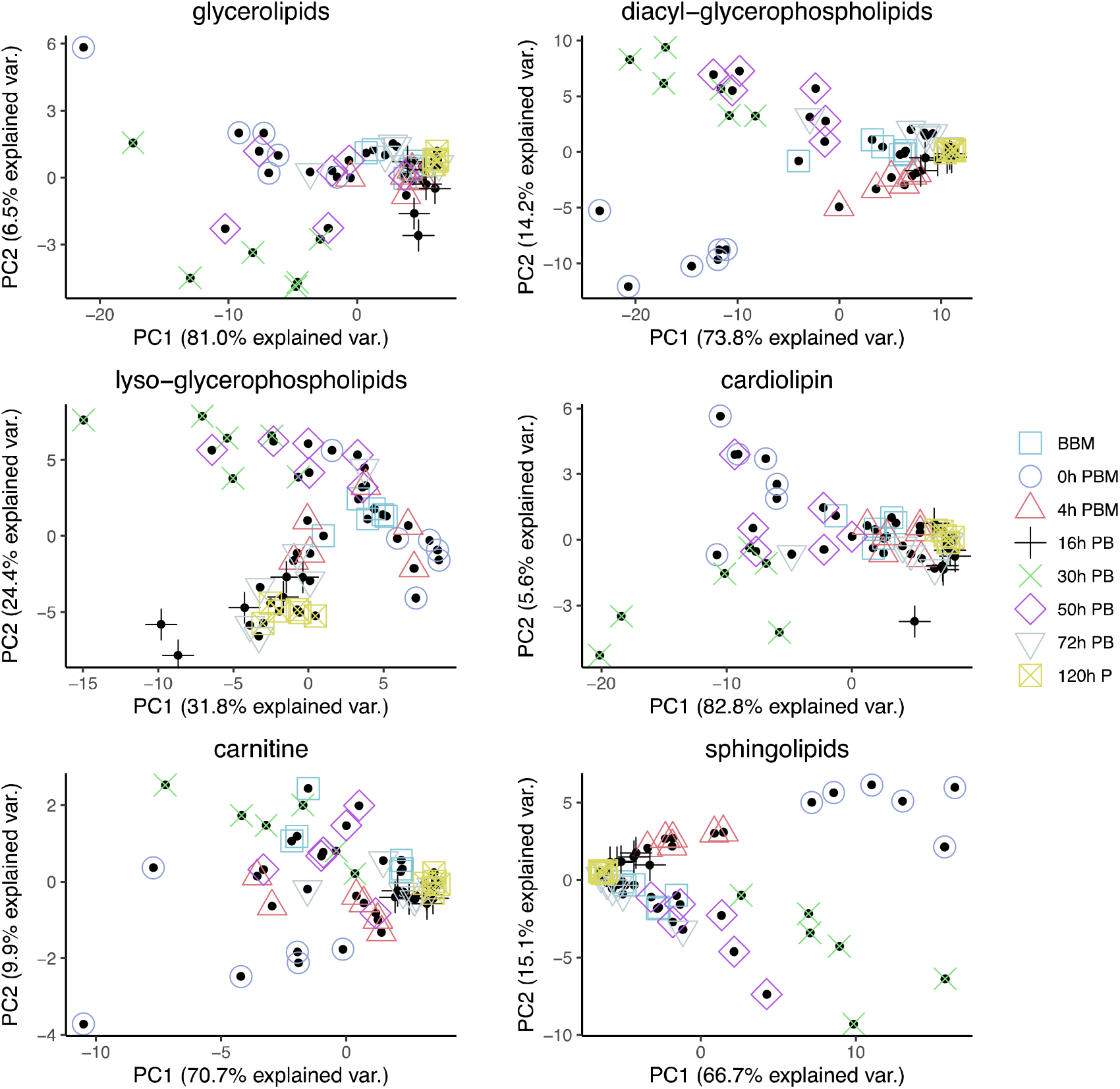
Principal Component Analysis of different groups of lipids in *Aedes aegypti* females at different time points over the course of feeding and reproduction. Colour and shape symbols correspond to the eight time points before blood meal (BBM) and post blood meal (PBM).

### Identification of change patterns for specific lipids

The amount of blood a female mosquito takes in when fully engorged can be similar to the weight of the individual, or even nearly twice the individual’s weight [35,36]. By comparing the time points BBM and 0 h PBM for individual lipid, we were able to determine which lipids were taken in with human blood. We found 86% (392 molecules) of all the lipids identified significantly increased at the time of blood meal (Supplementary Material 2). An exception occurred for glycerophospholipids and carnitines. We found 29 lyso-glycerophospholipids and 19 diacyl-glycerophospholipids that did not increase significantly after the blood meal, which might indicate that these lipids originated from mosquitoes instead of the human blood. These lipids include most of the LPA, LPG identified, and all LPI identified. The carnitines showed a mild increase generally. The carnitines tetradecasphingenine (C14) and hexadecasphingenine (C16) CAR did not change significantly, while most of the other carnitines (7 out of 11) increased only slightly (adjusted 0.05 > p > 0.01); only one molecule (CAR 20:1) in this class increased highly significantly (adjusted p < 0.001).

From 0 h to 16 h PBM, the amount of lipids dropped while the weight of the mosquitoes did not decrease correspondingly. From 16 h to 30 h PBM, 87.7% of lipids (400 molecules) identified increased significantly (Supplementary Material 3), of which 392 showed at least a two-fold increase, and 139 showed an increase of at least ten-fold. There were 34 lyso-glycerophospholipids which did not increase significantly and these lipids were distributed in all sub-classes of lyso-glycerophospholipids except for LPA. All LPA identified increased more than ten-fold, while none of the individual lipids identified from LPC and LPE significantly increased.

Unlike lyso-glycerophospholipids, almost all diacyl-glycerophospholipids had significantly increased, including all ether PI, oxidized PE, PE, PA, PG, PI and PS, In particular, 30 out of 43 PE and 18 out of 33 PI identified increased more than ten-fold. Besides, 42 out of 66 CL increased more than ten-fold and the four HexCer identified increased about ten-fold. In summary, lipids that increased by more than ten-fold include most of PE, PI and CL that act as membrane components, and all of HexCer and LPA that act as signalling molecules and participate in a variety of physiological functions. These lipids are expected to be particularly important in mosquito egg production.

In general, most individual lipids have similar change patterns, regardless of the length of fatty acyl chain and number of double bonds. Outliers (that did not increase significantly from BBM to 0 h PBM or from 16 to 30 h PBM) were found mostly in lyso-glycerophospholipids. Looking at the overall pattern, 21 lipid molecules identified did not increase from BBM to 0 h PBM or from 16 h to 30 h PBM, and 19 of these are lyso-glycerophospholipids.

We then further analysed the change patterns of lyso-glycerophospholipids, which represent a group composed of various sub-groups of lipids with different patterns of change (Figure 5), and diverse metabolic roles such as metabolic intermediates, extracellular signalling molecules and immune system activators. Specifically, the two most abundant lyso-glycerophospholipids LPE and LPC, which play important roles in membrane composition, were relatively stable in terms of total amount during reproduction. LPA, an important group of lipids in regulating various signalling pathways, showed a dramatic increase from 16 to 30 h PBM, which is similar to the change patterns for other groups (other than lyso-glycerophospholipids) described previously. LPI, LPG and LPS, on the other hand, only had one significant increase at a time period from 0 to 16 h PBM, when most lipids in other groups decreased. LPI, a bioactive lipid, can initiate a variety of functions such as cell growth, differentiation, and immunity [37,38]. While LPS and LPG, membrane lipids in gram-positive bacteria and gram-negative bacteria respectively, can act as microbial toxins to trigger mosquito immune response by activating the Toll and Imd pathway [39,40]. In this study, the changes of these two lipid groups most likely indicate participation of gut bacteria in mosquito blood digestion.

**Figure 5.**
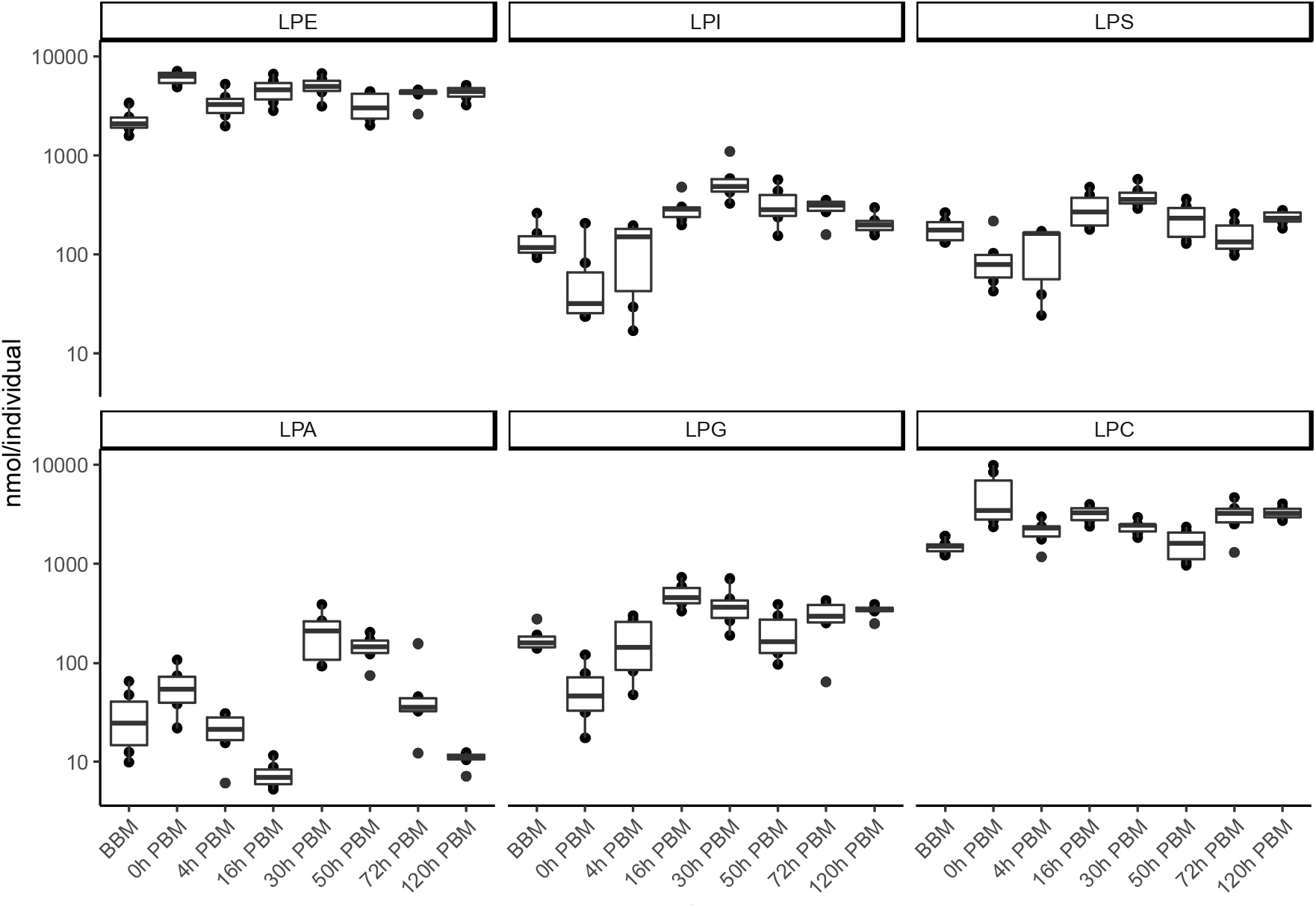
Changes of lipid amounts in lyso-glycerophospholipids in *Aedes aegypti* females over the course of feeding and reproduction. The total amount (nmol/individual) of LPE; LPI; LPS; LPA; LPG and LPC were measured at eight time points before blood meal (BBM) and post blood meal (PBM).

By comparing the differences between the two periods (from BBM to 0 h PBM and from 16 h to 30 h PBM) when most increases occurred, we found that most lipids we identified were present in both human blood and mosquitoes, and also being affected by mosquito ovarian follicle development. Nevertheless, some lipids seemed to be present at a stable level in mosquitoes and were not significantly increased after blood feeding, such as tetradecasphingenine (C14) and hexadecasphingenine (C16) CAR, and most of the LPGs.

Some lipids seemed to exist at a much higher abundance in human blood but were not used in mosquitoes to support egg development, which we define here as those that increased more than ten-fold after blood feeding but did not increase significantly (adjust p > 0.05) at the later period. These include some Oxidized PE and sphingomyelin.

## Discussion

In the current study, we focused on lipidomic changes of whole female mosquitoes during the reproductive process. Whole-body investigations provide an overall picture of lipid changes that might not be evident when focusing only on organs or cells. We found biphasic increases in more than 80% of lipids identified at the time of feeding and from 16 to 30 h PBM. The amount of lipids dropped between these periods of increase, as also observed in the fat body [41]. The weight of the mosquitoes did not decrease at the first time point when lipids decreased, suggesting that lipids were transformed into different metabolites. Many metabolites can be produced within this period such as glycogen, and fatty acids [20,42], but it is more likely that the decrease of lipids from 0 to 16 h PBM reflects the synthesis of vitellogenin in the fat body, which is triggered by blood feeding. Previous studies found that the secretion of vitellogenin in the fat body reaches its highest peak at around 18 h PBM, and decreases to the previtellogenic level at around 30 h PBM [43,44].

Pitch et al. [41] observed a relatively lower level of TG in the *Ae. aegypti* female fat body at around 30 h PBM, while our study of whole body lipidome showed highest level of TG at 30 h PBM. The differences may reflect the transportation of lipids from the fat body or midgut into oocytes during the insect reproductive process through lipophorin, a major lipid-carrying protein [45]. *Ae. aegypti* and some other Diptera predominantly transport TG via lipophorin, not DG as is the case of most other insects [46–48]. Moreover, a general observation for insects is that the midgut can convert DG to TG in the presence of high concentrations of potentially toxic fatty acids; this conversion could maintain low intracellular levels of DG and fatty acids to prevent lipotoxicity [20,49,50]. Additionally, there are other possibilities for lipid movement not based on DG transport including evolutionarily conserved mechanisms for movement of fatty acids [51].

Lyso-glycerophospholipids were highlighted as a lipid group of interest by the present study due to their unusual temporal pattern in abundance. Lyso-glycerophospholipids have diverse functions and they are also found in cell membranes [52]; as precursor of phospholipid biosynthesis, they may act as signal molecules involved in a broad range of biological processes [53,54]. However, they might also reflect a highly regulated reservoir of lipids with accessible utility and concentration-dependent lipid toxicity liability [55], which can result from lipid overload generated by dietary excess [56] as might be the case when mosquitoes blood feed. LPE and LPC incorporate into membranes and change membrane permeability and stability. The change in permeability and stability can result in membrane toxicity unless concentrations of lyso-glycerophospholipids are kept below a cytotoxic limit [55], which may explain the stability of these two lipid groups. Also, we found substantial increases in cell membrane lipids between 16 and 30 h PBM as in the cases of PE and PC, reflecting the growth of embryos during the reproductive process. Moreover, lyso-glycerophospholipids could act as a collective reserve which provides acyl chains for beta-oxidation when needed [51], or they could be a source of partially assembled molecules for further biosynthesis, which may explain the unusual increase of LPI between 0 and 16 h PBM followed by increase of nearly all major lipid groups from 16 to 30 h PBM. LPS and LPG also increase between 0 and 16 h PBM, which might indicate the growth of gut bacteria during the blood digestion process [57]. LPA is the only group from lyso-glycerophospholipids showing a bimodal pattern with increases, and these lipids are essential signalling molecules [58].

We also found that many lipids from groups PI and CL increased dramatically from 16 to 30 h PBM. These lipids are closely connected with metabolic activity and signalling. For example, PIs are important in regulating phototransduction, cell growth and the developmental process of fruit flies [59–61]. And CLs are integral to mitochondrial membranes, with important roles in ATP-synthase and phospholipid remodelling [62,63]. Moreover, SM play significant roles in mammal cells being involved in diverse signalling pathways [64,65], whereas for dipteran insects such as *Drosophila*, there is a lack of SM and instead they synthesize the SM analogue phosphoethanolamine ceramide [66]. Though we did not look into phosphoethanolamine ceramide in the present study, we also note that most SMs were either relatively constant or changed only slightly between 16 h and 30 h PBM.

Finally, it is worth noting that approximately a quarter of the lipids, including almost all SHexCer we identified, contained odd chains. Pinch et al. [41] investigated the fat body of mosquitoes feeding on bovine blood and considered that odd-chain lipids may come from ruminal microflora or microbes in larval rearing water or branched chain amino acids from haemoglobin. Our study showed that the last hypothesis is most likely correct, as we observed these odd-chain lipids from human blood and these also increased in mosquitoes between 16 h and 30 h PBM. These odd chain fatty acid-containing lipids did not show any distinct change patterns comparable to even chain fatty acid-containing lipids from the same lipid group.

## Supplementary Materials

Supplementary Material 1. Ovarian appearance of female mosquitoes dissected at (A) BBM, (B) 1 d PBM, (C) 2 d PBM, (D) 3 d PBM, (E) 4 d PBM and (F) 5 d PBM. There is some variation in the stages of development of the ovaries among individuals as captured by variation in weight (Fig. 1).

Supplementary Material 2. List of individual lipids and their changes before and after female *Aedes aegypti* mosquitoes blood fed (between BBM and 0 h PBM).

Supplementary Material 3. List of individual lipids and their changes between 16 hours and 30 hours after female *Aedes aegypti* mosquitoes blood fed (between 16 h and 30 h PBM).

## Ethics statement

Female mosquitoes were blood fed on a volunteer in this experiment, with ethics approval from the University of Melbourne Human Ethics committee (approval 0723847).

## Acknowledgment

We thank Cameron Patrick from Statistical Consulting Centre and Melbourne Statistical Consulting Platform, the University of Melbourne, for statistical advice. We also thank Perran A. Ross and Moshe Jasper for their assistance and advice.

